# The lost vultures of Romania: reconstructing two centuries of decline from historical records and citizen science data (*Gyps fulvus*, *Aegypius monachus*, *Neophron percnopterus, Gypaetus barbatus*)

**DOI:** 10.64898/2026.05.13.723308

**Authors:** Gergely Osváth, Avar-Lehel Dénes, Zsolt Kovács, Alexandru-Cătălin Birău, Edgár Papp, Gabriella-Veronka Jákó, Róbert Zeitz

**Affiliations:** Museum of Zoology, Babeș-Bolyai University, Cluj-Napoca, Romania; Evolutionary Ecology Group, Babeș-Bolyai University, Cluj-Napoca, Romania; STAR-UBB Institute for Advanced Studies in Science and Technology, Babeș-Bolyai University, Cluj-Napoca, Romania; Faculty of Biology and Geology, Babeș-Bolyai University, Cluj-Napoca, Romania; Văcărești Natural Park Administration, Bucharest, Romania; Milvus Group-Bird and Nature Protection Association, Târgu Mureș, Romania; Foundation Conservation Carpathia, Brașov, Romania

**Keywords:** Griffon Vulture, Cinereous Vulture, Egyptian Vulture, Bearded Vulture, Carpathians, Dobrogea, historical ornithology, museum collections, reintroduction, population decline

## Abstract

Romania represents one of the few European Union member states in which all four Old World vulture species historically maintained breeding populations: the Griffon Vulture (*Gyps fulvus*), Cinereous Vulture (*Aegypius monachus*), Egyptian Vulture (*Neophron percnopterus*) and Bearded Vulture (*Gypaetus barbatus*). Until ongoing reintroduction efforts, Romania remained among the last EU countries whose former vulture guild had not yet been targeted for active recovery. Despite this exceptional significance in a European conservation context, no comprehensive synthesis of the historical and contemporary distribution of these species in Romania has been undertaken. We conducted an extensive review to gather all available vulture occurrence data and present a database of 1,136 occurrence records spanning 1818–2025. We systematically searched museum collections, historical ornithological literature, modern field surveys and citizen-science platforms. The database documents substantial breeding populations across the Carpathian arc and Dobrogea until the early twentieth century, followed by near-total breeding collapse between the 1920s and 1960s driven by persecution, secondary poisoning and agrarian transformation. In total, 148 confirmed or probable breeding records have been documented for the four species combined, with the most recent confirmed breeding records dating to 1929 (*Gyps fulvus* and *Gypaetus barbatus*), 1942 (*Aegypius monachus*) and 1966 (*Neophron percnopterus*). Non-breeding occurrences increase markedly after 2010, consistent with dispersal from expanding Balkan source populations. The Făgăraș and Retezat Mountains emerge as the historically most important breeding strongholds for three of the four species (all except the Egyptian Vulture). This dataset highlights the value of reconstructing historical baselines where functional extinction preceded modern monitoring, and provides an empirical foundation for reassembling a keystone scavenger guild in the Carpathian–Danube region.

## 1. Introduction

Old World vultures fulfil irreplaceable ecological roles as obligate scavengers, accelerating nutrient cycling and reducing pathogen transmission through carcass removal (Donázar et al. 2016, Moleón et al. 2014). European vulture populations have nonetheless declined severely over the past two centuries through direct persecution, secondary poisoning by veterinary pharmaceuticals and pesticides, infrastructure collisions, food shortages from livestock-husbandry change, and loss of cliff-nesting habitat (Botha et al. 2017, BirdLife International 2024). By the mid-twentieth century, breeding populations across most of Central and Eastern Europe had been extirpated, leaving the Iberian Peninsula, the Balkans and a few Mediterranean islands as the principal remaining strongholds (Ogada et al. 2012). Globally, 14 of the 23 recognised vulture species are currently classified as Threatened or Near Threatened (Ogada et al. 2012, BirdLife International 2024).

In countries where vultures disappeared before the advent of systematic monitoring, the former presence and ecological significance of these species is often poorly documented (shifting-baseline syndrome; Pauly 1995); without explicit reconstruction of historical baselines, conservation targets risk being set too low and the spatial extent of formerly suitable habitat underestimated. Targeted programmes have, however, demonstrated that recovery is achievable: the Bulgarian Rhodopes Griffon Vulture population has grown from c. 20 pairs in the 1970s to over 163 pairs today (Terraube et al. 2022), and Griffon Vulture reintroductions in Sardinia, France, Bulgaria, Cyprus and Croatia have established new populations across the species’ former range. The Bearded Vulture has been restored to the European Alps, with the metapopulation now exceeding 70 territorial pairs and producing dispersers reaching the Balkans and Scandinavia (Frey et al. 2022). Increasing vagrancy of the Griffon Vulture into Central Europe has been documented since the early 2000s, with ringing recoveries confirming Balkan origin (Danko et al. 2013); a recent monograph by Tóth (2022) provides a parallel synthesis for Hungary and the Carpathian Basin.

Within this broader European context, Romania occupies a position of extraordinary importance for vulture conservation – and, arguably, for European obligate scavenger ecology more broadly. It is one of the few current EU member states in which all four Old World vulture species – Griffon Vulture, Cinereous Vulture, Egyptian Vulture and Bearded Vulture – formerly maintained confirmed breeding populations (Dombrowski 1912, Tălpeanu 1967, Papadopol and Tălpeanu 1986). Historical sources paint a vivid picture of former abundance: Landbeck (1842, cited in Salmen 1980) described the Griffon Vulture as “the most common vulture in Transylvania”; Crown Prince Rudolf called it “the most common bird of the Transylvanian Alps” in the early 1880s (Salmen 1980); and Dombrowski (1912) counted 52 Griffon Vultures on a single oak tree in the Ghiuvegea forest of Dobrogea, estimating at least 300 individuals roosting there on one evening. For the Cinereous Vulture, the Sintenis brothers collected 377 eggs from the Babadag forest alone during 1873–1875, implying a breeding colony of well over 100 pairs (Dombrowski 1912). Ádám Buda reported the Bearded Vulture as “very common” in the Transylvanian mountains in the early 1890s, where he daily saw 2–3 and sometimes over 10 at once (Buda 1906). The Carpathian Mountain arc, with its extensive montane cliff systems, sub-alpine grasslands and traditional transhumant livestock economy, and the Dobrogean plateau, with its steppic character, rocky gorges and proximity to the major Black Sea flyway, historically supported the complete European obligate scavenger guild – the four-species assemblage whose simultaneous presence is today found only in parts of the Iberian Peninsula and the Balkans. Nineteenth-century ornithologists documented colonies of Griffon and Cinereous Vultures in the Retezat and Rodna Mountains rivalling those of contemporary Balkan strongholds, while the Bearded Vulture bred in the Făgăraș, Retezat and Apuseni ranges until well into the twentieth century.

The progressive disappearance of all four species from Romania is documented in regional ornithological accounts (Csató 1885, Frivaldszky 1891, Dombrowski 1912, Schenk 1918, Salmen 1980, Papadopol and Tălpeanu 1986) scattered across multiple languages (German, Hungarian, Romanian, English and French), but no synthesis integrating museum specimens, the large body of historical literature and modern citizen-science records has previously been assembled. Such a synthesis is increasingly recognised as a prerequisite for evidence-based reintroduction planning (Seddon et al. 2014, Armstrong and Seddon 2008): without an established historical baseline, defining recovery targets, selecting release sites and evaluating progress against pre-decline parameters become difficult.

The study has three main objectives: (1) to compile the most complete database of vulture occurrence records in Romania currently achievable, including all available records up to 31 December 2025, by integrating museum collections, published and unpublished historical literature, and citizen science data; (2) to describe the temporal and spatial patterns of occurrence for all four species, with particular attention to breeding localities, the chronology of population decline and the scale of recent vagrant movements; and (3) to identify the key historical sites and habitats of conservation significance and propose areas that merit consideration for future reintroduction efforts based on the strength of the historical evidence.

## 2. Methods

We compiled all available occurrence data for the target species (*Gyps fulvus*, *Aegypius monachus*, *Neophron percnopterus*, *Gypaetus barbatus*) within the territory of present-day Romania. The search was unrestricted backward in time, with an inclusion cutoff of 31 December 2025; the earliest record successfully extracted dates to 1818. For the full list of sources, detailed methodology and the complete reference list of occurrence records, see Supplementary Material S1; the complete occurrence database is provided in Supplementary Material S2.

The dataset was assembled through systematic searches for vulture occurrence data across four categories of source: (i) published ornithological literature, including the major historical regional syntheses (e.g., Frivaldszky 1891, Csató 1885, Bielz 1888, Hausmann 1877, Tschusi zu Schmidhoffen 1888, Spiess 1898, Dombrowski 1912, Schenk 1918, Salmen 1980, and Tóth 2022, drawing on the archives of the Hungarian Ornithological Centre and the journal Aquila, which constituted the single largest contributing source with 162 individually attributable records), publications and museum catalogues (e.g., Cătuneanu et al. 1967, Tălpeanu 1967, Cătuneanu 1973, Papadopol and Tălpeanu 1986, Osváth et al. 2022, Mestecăneanu 2021), and the regional hunting and natural history journals (systematically searched); numerous additional works were consulted but did not yield further vulture records, and only credible records were retained, with selection criteria specified in Supplementary Material S1 (which also lists all publications consulted, including non-yielding sources); (ii) specimen and archival data from Romanian natural history collections (Supplementary Material S1); (iii) records from citizen-science platforms (Rombird, Ornitodata, openbirdmaps, eBird) — for citizen-science records, only the first observation of each identifiable individual was retained to avoid pseudoreplication of repeat sightings; and (iv) unpublished field data contributed by the authors, including standardised monitoring surveys in the Făgăraș Mountains. Records were categorised as historical (pre-2000) or recent (2000 onwards), following Birău et al. (2024).

Where secondary or tertiary compilations referenced earlier observations, we identified and cited the original primary publication directly (e.g., Hausmann 1877; Tschusi zu Schmidhoffen 1888; Spiess 1898); the regional compilation served only to identify the primary source.

Each record was assigned values for species, locality, county, historical region, decimal coordinates, date, year, number of individuals, record type, status, observer, age, sex, source citation and notes. Coordinates were assigned by reference to the original locality descriptions, supported by historical maps and contemporary 1:50,000 topographic maps; locality precision was classified into four ordinal categories: exact (≤ 1 km, e.g. specific named point such as a peak, village centre, or museum specimen with locality detail); approx (1–30 km, e.g. identifiable named area such as a single mountain range, valley, or commune); region (≥ 30 km, e.g. county-level, multi-range mountain system, historic region, or national-scale reference); and GPS (5–50 m, modern citizen-science platform GPS-derived coordinates). For records described only at landscape scale or larger, centroid coordinates were used with the corresponding precision flag. All records retain coordinates regardless of precision class to enable region-scale visualisation; for fine-grained spatial analyses, region-precision records can be filtered out. Where secondary compilations conflicted with primary sources, the primary source took precedence. Undated records attributed to a named observer were assigned a terminus ante quem based on the observer’s active period rather than the publication date. Duplicate records (the same physical observation cited independently in multiple compilations) were identified through a two-stage review of 203 potential duplicate groups (521 candidate records), and 258 confirmed duplicates were removed, with unique information merged into the retained record. Full quality-control protocols — duplicate detection, taxonomic standardisation, temporal consistency checks and geographic validation — are described in Supplementary Material S1. For records whose source described a date range rather than a single year, the Year field was set to the mid-range estimate, with the original range preserved in the Date field; records with date uncertainty exceeding ∼20 years (n=3) are flagged in Notes for caution in decade-level analyses.

Records were summarised by species, decade, biogeographic region and county. Breeding records – confirmed or probable nesting evidence (nest, egg, chick, or unambiguous breeding behaviour) – were analysed separately to trace the chronology of population decline. Each breeding record was classified using the European Breeding Bird Atlas evidence categories (Keller et al. 2020): Confirmed (C: nest with eggs or young, recently fledged young attended by adults, or directly observed occupied nest), Probable (B: nest reported without direct chick or egg detail, territorial pair, nest building, or pair in suitable habitat in the breeding season), and Possible (A: single bird in suitable habitat during the breeding season, or vague historical regional claim). EBBA evidence categorises the biological observation; the Reliability column (Certain / Probable / Uncertain / Not assessed) records the strength of source documentation. The last EBBA Confirmed breeding record per species was used to define the breeding endpoint. Records lacking an inferable year (n = 101) were excluded from temporal analyses but retained in the database. Spatial analysis and mapping were performed using QGIS 3.40.1. All data are available in Supplementary Material S2.

## 3. Results

### 3.1 Overview of the database

The final database comprises 1,136 records (1,036 georeferenced) of four vulture species from Romania, spanning 1818-2025 (Table 1). The breakdown by species is: Cinereous Vulture n = 342 (30.1%); Griffon Vulture n = 306 (26.9%); Bearded Vulture n = 315 (27.7%); Egyptian Vulture n = 173 (15.2%). The database contains 148 confirmed or probable breeding records across all species combined (Table 2; Figure 1). Specimen records (mounted specimens, bird skins and skeletal material) number 209 in total, documented through direct natural history collection visits, published specimen catalogues (Table 3) and historical literature.

**Table 1.**
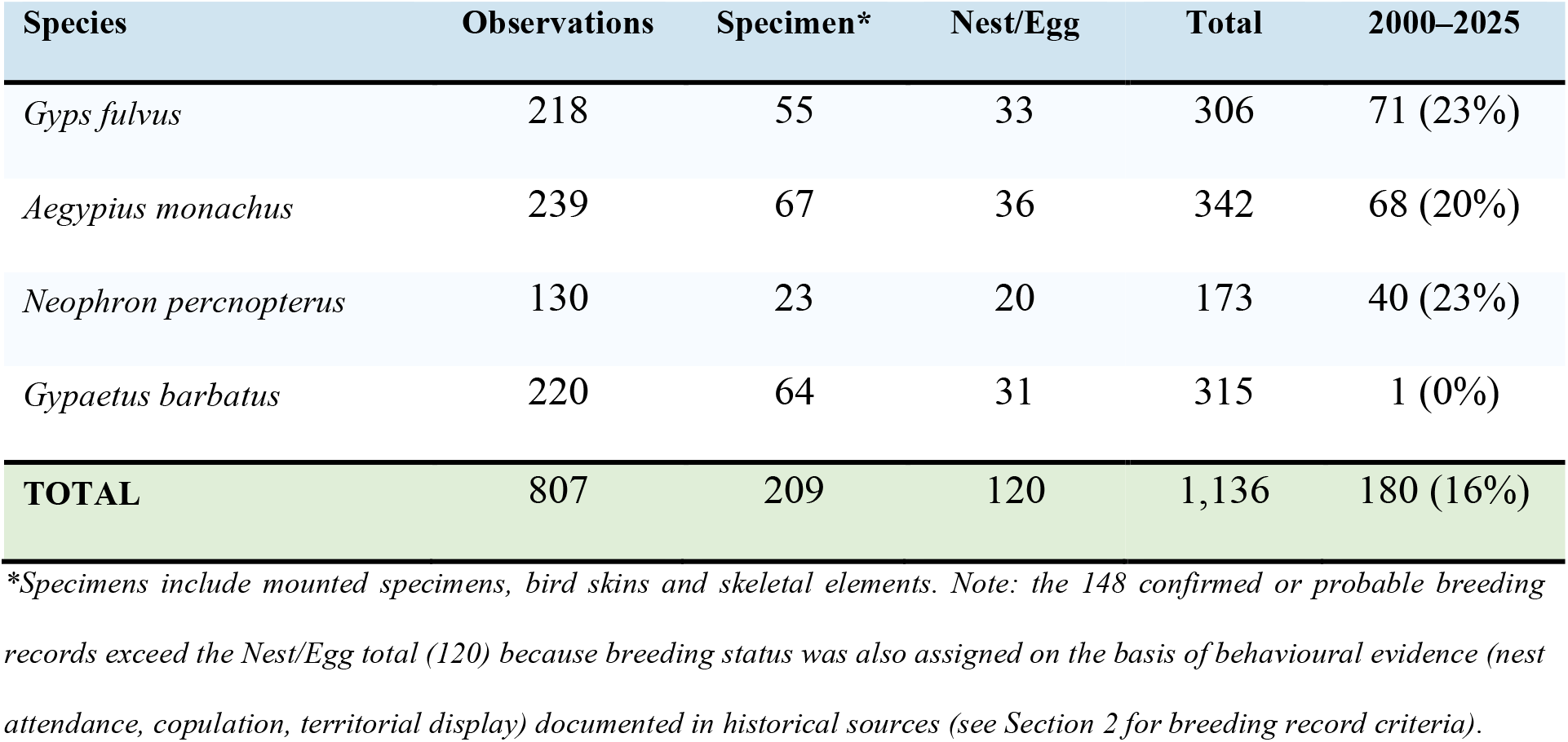
Summary of the vulture occurrence database by species and record category.

| Species | Observations | Specimen* | Nest/Egg | Total | 2000–2025 |
| --- | --- | --- | --- | --- | --- |
| *Gyps fulvus* | 218 | 55 | 33 | 306 | 71 (23%) |
| *Aegypius monachus* | 239 | 67 | 36 | 342 | 68 (20%) |
| *Neophron percnopterus* | 130 | 23 | 20 | 173 | 40 (23%) |
| *Gypaetus barbatus* | 220 | 64 | 31 | 315 | 1 (0%) |
| **TOTAL** | **807** | **209** | **120** | **1,136** | **180 (16%)** |

\*Specimens include mounted specimens, bird skins and skeletal elements. Note: the 148 confirmed or probable breeding records exceed the Nest/Egg total (120) because breeding status was also assigned on the basis of behavioural evidence (nest attendance, copulation, territorial display) documented in historical sources (see Section 2 for breeding record criteria).

**Table 2.**
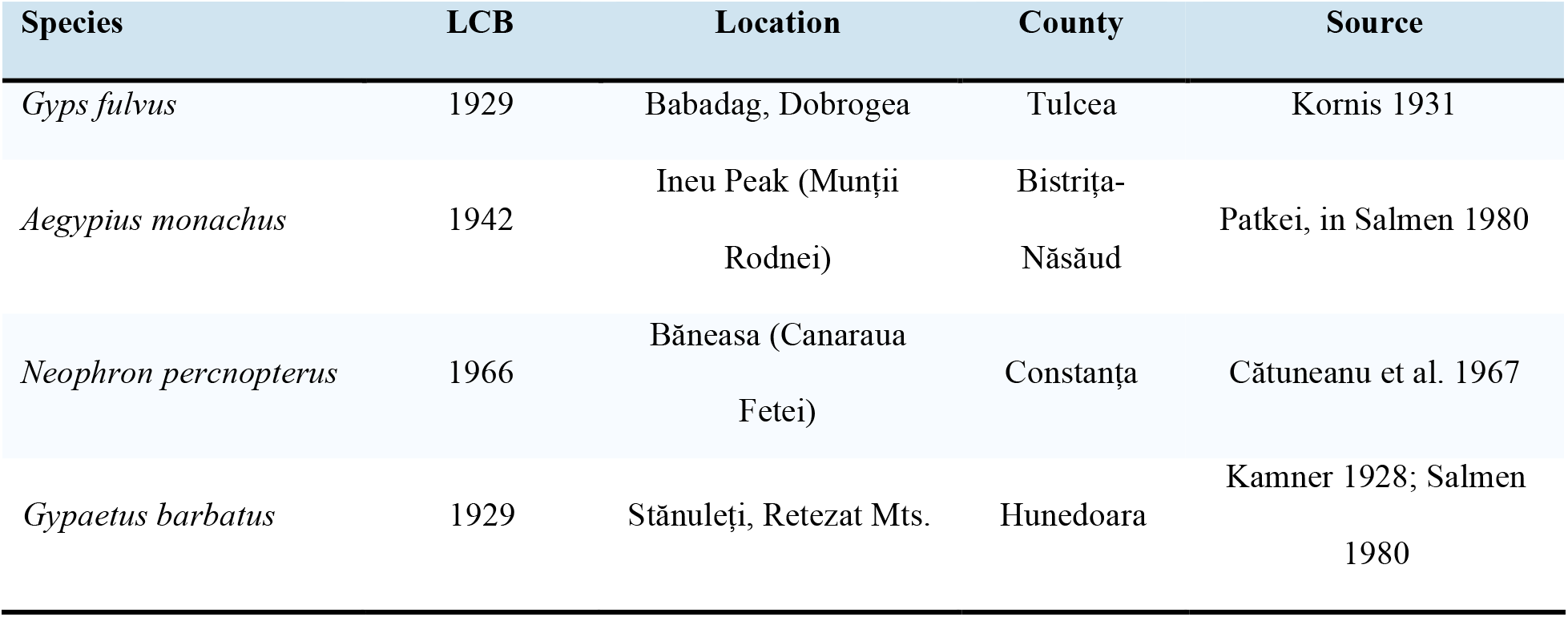
Last confirmed breeding (LCB) records for each vulture species in Romania.

**Table 3.** Data sources by category and temporal period.

| Source category | Records (n) | % of total | Definition |
| --- | --- | --- | --- |
| *Historical literature (pre-1950)* | 783 | 68.9% | Pre-1950 published records (literature, treatises, regional faunas) |
| *Museum collections* | 108 | 9.5% | Romanian natural history museum specimens (catalogues + direct visits) |
| *Modern field literature (1950+)* | 78 | 6.9% | Post-1950 Romanian published field surveys and authors' unpublished field data |
| *Citizen-science platforms* | 167 | 14.7% | Rombird, Ornitodata, openbirdmaps, eBird, iNaturalist (mostly 2010+) |
| **Total** | **1,136** | **100%** |  |

**Figure 1.**
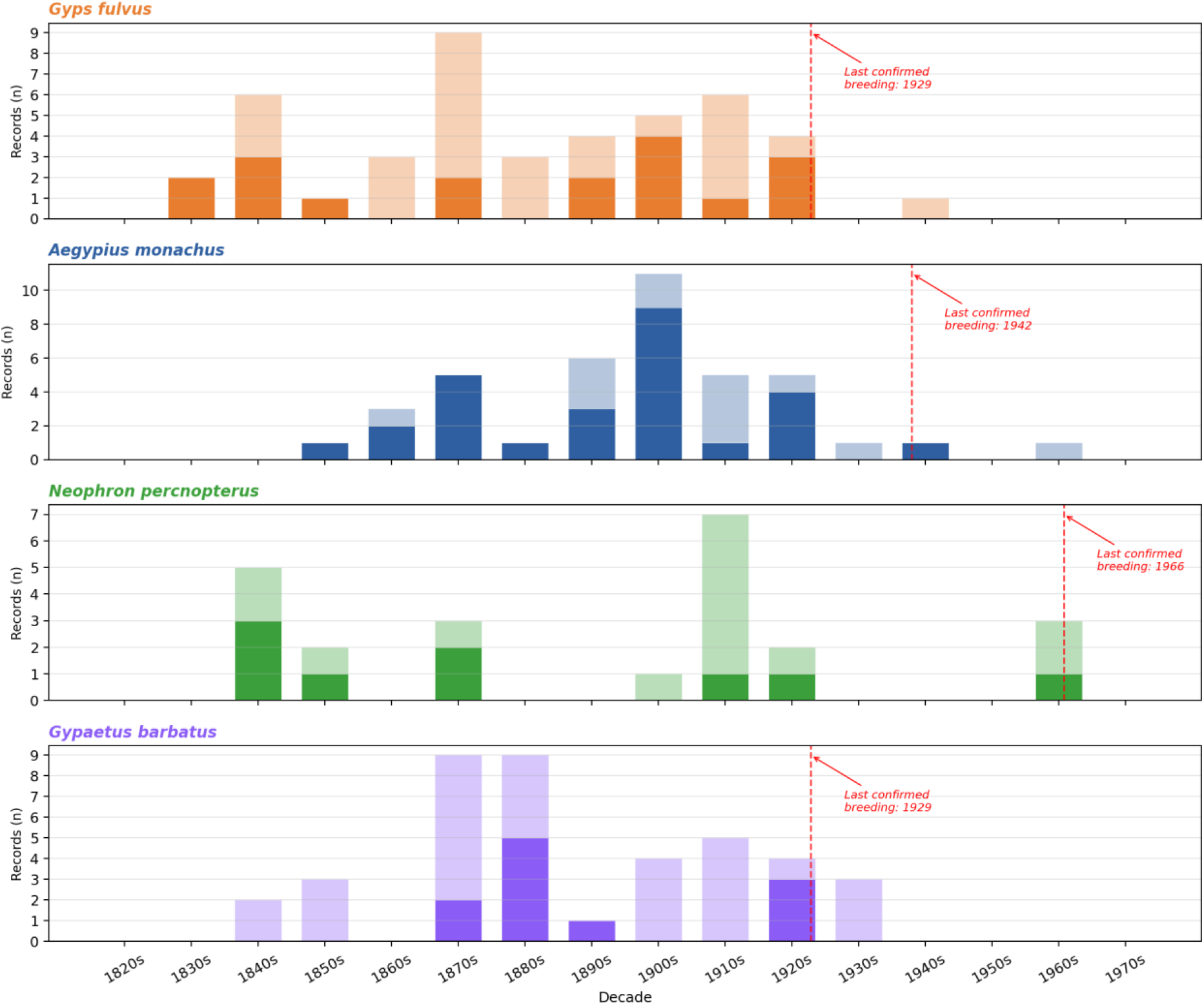
Breeding records per decade for each species, 1820–1975. Stacked bars: solid colour = EBBA Confirmed breeding (category C: nest with eggs or young, fledged young with adults, or directly observed occupied nest); paler shade = EBBA Probable or Possible (categories B and A: nest reported without direct chick or egg evidence, territorial pair, or pair / single bird at a suitable site during the breeding season).

The largest source category is historical published literature (pre-1950 Transylvanian, Hungarian and German ornithological works and their modern compilations), contributing 783 records (68.9%). Museum collection specimens documented through published catalogues or direct collection visits contribute 108 records (9.5%); post-1950 Romanian published field literature together with unpublished field data contributed by the authors comprise 78 records (6.9%); and citizen-science platforms (Rombird, Ornitodata, OpenBirdMaps, eBird, iNaturalist) contribute 167 records (14.7%), predominantly from 2010–2025 (Table 3).

Geographically, Transylvania accounts for the largest share of records (n = 653; 57.5%), followed by Dobrogea (n = 196; 17.3%), Banat (n = 106; 9.3%), Muntenia (n = 72; 6.3%), Oltenia (n = 53; 4.7%) and Moldova (n = 19; 1.7%; Figure 2). The resulting dataset represents heterogeneous historical observations with highly variable detectability across periods and regions; all temporal and spatial patterns described below should be interpreted accordingly. The sources of this geographic and temporal bias are addressed in the Discussion. At county level, records concentrate in the southern Carpathian counties (Hunedoara, Sibiu, Brașov, Caraș-Severin) and northern Dobrogea (Tulcea, Constanța), reflecting the historical distribution of breeding populations and museum collection geography (see Figure 2).

**Figure 2.**
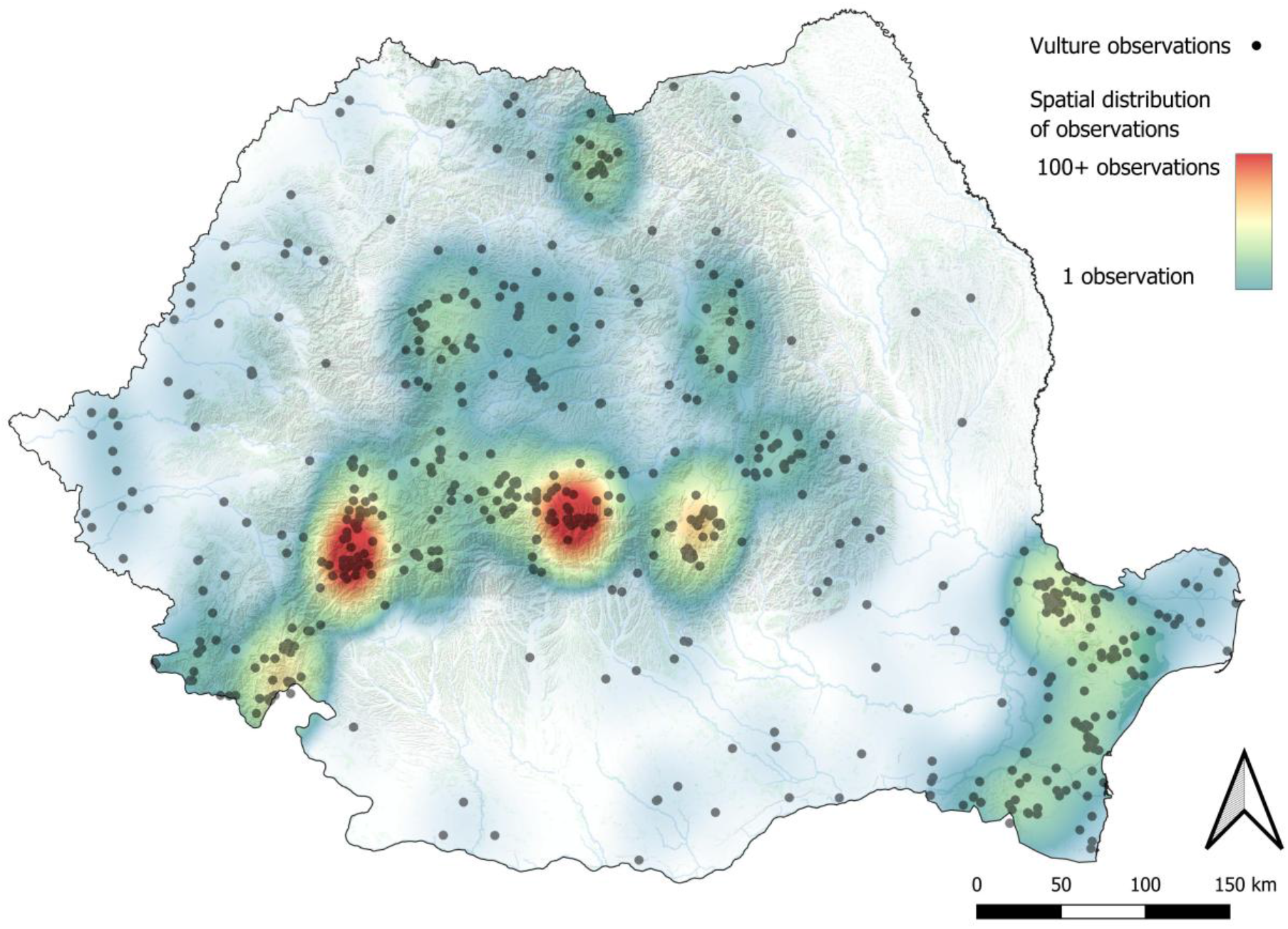
Spatial distribution of all georeferenced vulture occurrence records in Romania (n = 1,036, all four species combined). Black dots = individual records; coloured surface = kernel-density estimate of observation intensity (cool colours = sparse, warm colours = dense)

### 3.2 Temporal patterns

The number of records per decade reflects both genuine changes in vulture abundance and substantial variation in observer effort (Figure 3). Among historical decades, the 1880s show the highest record total (n = 146), coinciding with the peak of systematic ornithological activity in Transylvania and the publication of major regional avifaunas. Record numbers in the 1900s and 1910s remain relatively high (n = 108 and n = 126 respectively), before declining sharply from the 1930s as vulture populations collapsed and ornithological activity decreased. The 1940s–1980s are severely under-documented (n = 2 in the 1970s, n = 5 in the 1980s), reflecting both genuine scarcity and a near-complete absence of systematic fieldwork on these species during this period.

**Figure 3.**
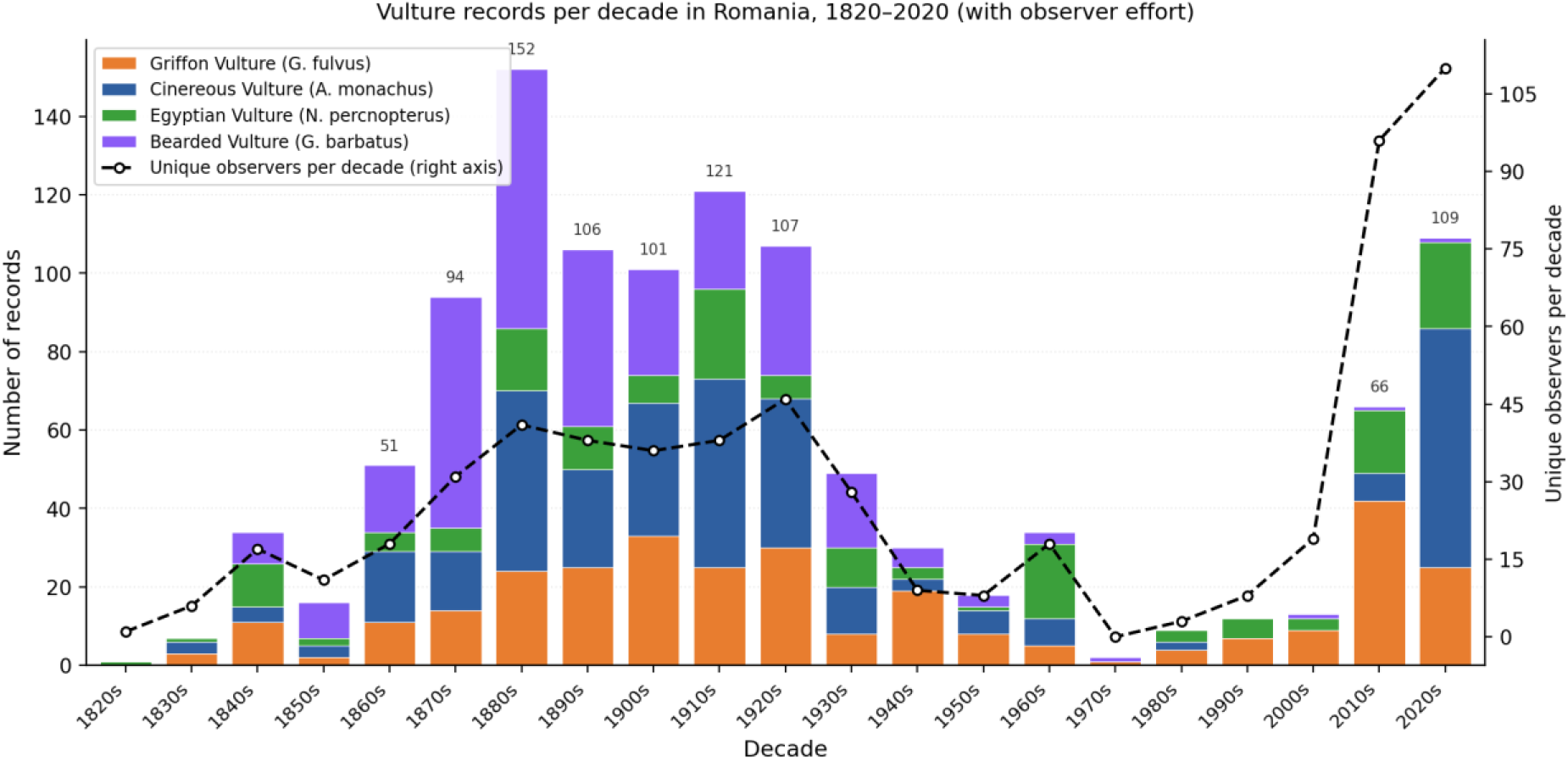
Number of vulture records per decade in Romania, 1818–2025, stacked by species (coloured bars). The black dashed line shows the number of unique observers per decade (right axis), used as a proxy for observer effort. The sharp increase from 2010 onward reflects the growth of citizen-science data; the near-absence of records in the 1940s–1980s reflects both genuine scarcity and very low observer effort during this period.

A marked recovery in records is evident from 2010 onward, driven entirely by citizen science: 63 records in 2010–2019 and 105 records in 2020–2025 alone (Figure 3). The 148 confirmed or probable breeding records are strongly concentrated in 1840–1930 (n = 135; 91%), with six records in 1931–1946 and four in 1957–1966 representing the final confirmed breeding attempts of any vulture species in Romania. Across all four species, the breeding record reveals a broadly synchronous pattern of guild-level collapse: the number of counties holding confirmed breeding records declined from a peak of at least 12 in the 1880s–1900s to three by the 1940s and zero after 1966.

### 3.3 *Gyps fulvus* (Griffon Vulture)

A total of 306 records of Gyps fulvus are included in the database, with 276 georeferenced records (90.2%; Supplementary Material S2; Figure 4). The earliest detailed account of breeding colonies is from Baldamus (1851), who found nesting groups at the Kazan Gorge and in the Czerna Valley, with nests on rock ledges, under overhangs and in caves, and observed flocks of 24 and 12 individuals soaring near the Danube. Dombrowski (1912) counted 52 Griffon Vultures on a single tree at the Ghiuvegea forest roost. The species has the longest documented history of occurrence in the dataset, with the earliest record from 1818 and the most recent from 2025. Records are concentrated in Tulcea (n = 41), Sibiu (n = 31), Constanța (n = 22), Hunedoara (n = 21) and Mehedinți (n = 19) counties.

**Figure 4.**
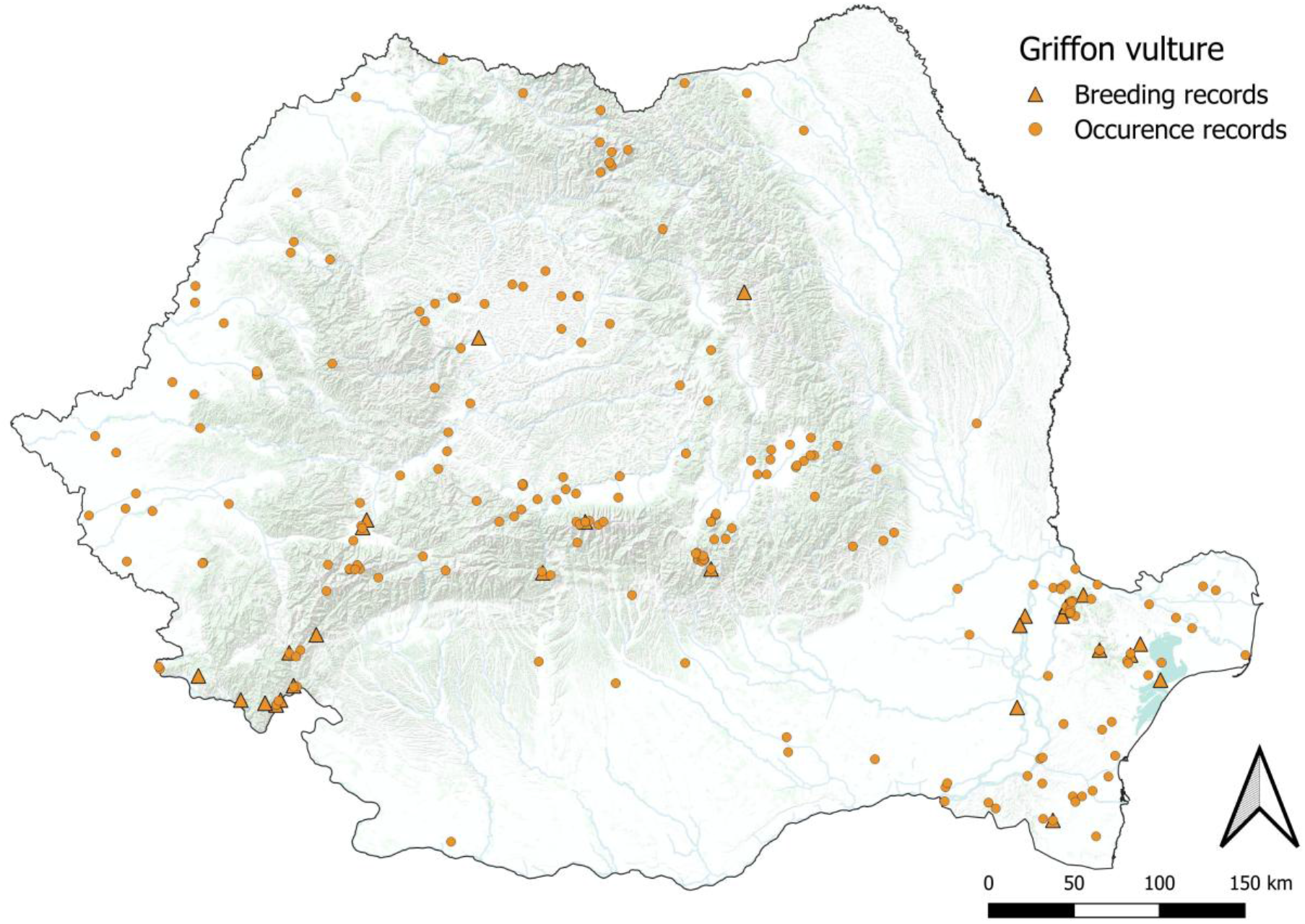
Distribution of *Gyps fulvus* (Griffon Vulture) in Romania. Orange circles = all occurrence records (n = 306); orange triangles = confirmed or probable breeding records (n = 44).

Forty-three confirmed or probable breeding records span 1835–1941. The most important historical breeding locality was the Cazane Gorge in Caraș-Severin County, where breeding is documented continuously from 1835 to 1923 (seven records). Dobrogea, particularly the Babadag Forest area in Tulcea County, yielded four breeding records, with the last confirmed breeding in Romania for this species in 1929 (Kornis 1931) at Babadag Forest (Dombrowski 1912, subsequent literature). Additional isolated breeding records come from Turda (Cluj, 1848), the Giurgeu Mountains (Harghita, 1860) and Sinaia (Prahova, 1862).

Non-breeding records show a strong bimodal temporal distribution: a peak in the 1880s– 1920s corresponding to the period of intensive ornithological documentation, a near-complete absence from the 1930s to the 2000s, and a substantial increase from 2010 onward (n = 62 records in 2010–2025).

### 3.4 *Aegypius monachus* (Cinereous Vulture)

With 342 records, Aegypius monachus is the best-represented species in the database, with 296 georeferenced records (86.5%; Supplementary Material S2; Figure 5). The temporal span is 1835–2025. Historical breeding records include remarkable nest descriptions: Spiess (1890, in Salmen 1980) described a nest 1.5 m in diameter on a lightning-broken beech tree in the Dobra valley, built from arm-thick branches; and a forest guard at Ghiuvegea reported a nest continuously occupied for 31 years, measuring 2.60 m across and 2.40 m high (Dombrowski 1912). The species shows the widest geographic distribution among the four target species in the historical record, with confirmed breeding in multiple distinct regions. County record totals are highest in Sibiu (n = 43), Hunedoara (n = 39), Tulcea (n = 35), Cluj (n = 27) and Constanța (n = 23).

**Figure 5.**
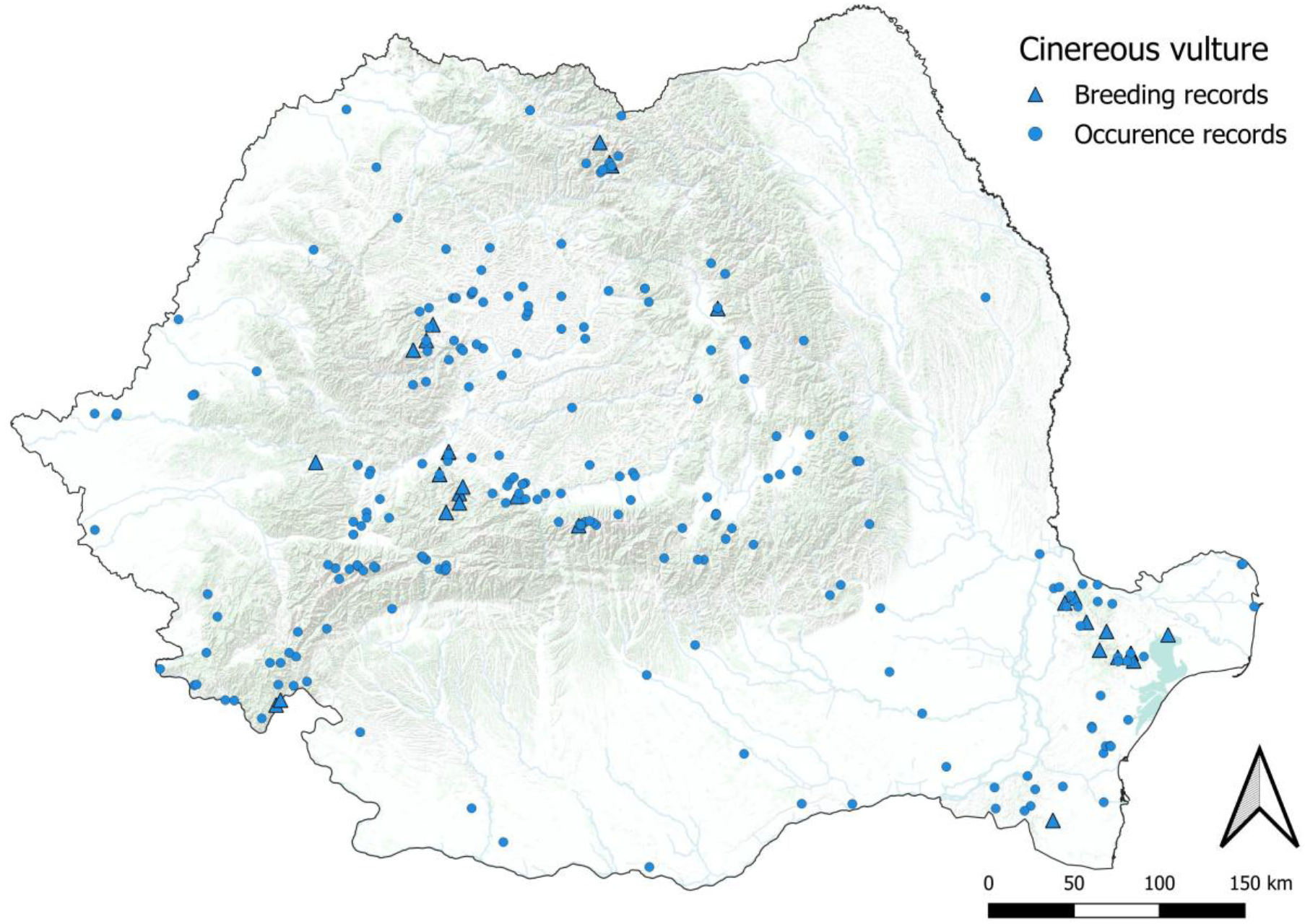
Distribution of *Aegypius monachus* (Cinereous Vulture) in Romania. Blue circles = all occurrence records (n = 342); blue triangles = confirmed or probable breeding records (n = 41).

Forty-one confirmed or probable breeding records span 1837–1964, with the last documented nest at Rodna (Ineu Peak) in May 1942 (Patkei, in Salmen 1980). The last breeding is reported as c. 1964 (Baumgart 1974a, citing Klemm 1966), while the last field observations were recorded in the Retezat Mountains in 1966 (Filipașcu 1967; Pușcariu 1967). The historically most important breeding counties are Tulcea (n = 15; Babadag, Cocoș Monastery and Greci forests of northern Dobrogea), Sibiu (n = 6; Valea Dobrei, Făgăraș and Sibiu surroundings), Alba (n = 4; Valea Dobrei / Munții Sebeșului), Bistrița-Năsăud (n = 3; Rodna Mountains), Cluj (n = 3; Valea Ierii and Gilău Mountains) and Constanța (n = 3; Bașpunar, Mircea Vodă and Ghiuvegea). The earliest breeding record is from Rodna in 1855. Breeding at the Dobra Valley (Hunedoara) is documented from 1876 to at least 1906; at Babadag (Tulcea) to 1929; and in the Rodna Mountains (Bistrița-Năsăud) until the definitive record from Ineu Peak in 1942, which represents the last geographically precise breeding record for this species in Romania. Papadopol and Tălpeanu (1986) cite an undated record of approximately 1964 as the probable final breeding, though its location is unspecified.

### 3.5 *Neophron percnopterus* (Egyptian Vulture)

A total of 173 records of Neophron percnopterus are included, with 161 georeferenced (93.1%; Supplementary Material S2; Figure 6). This species shows the most restricted historical distribution of the four, strongly concentrated in two geographic centres: Dobrogea (Constanța County, n = 39; Tulcea County, n = 30) and the Iron Gates region (Caraș-Severin, n = 22; Hunedoara, n = 20).

**Figure 6.**
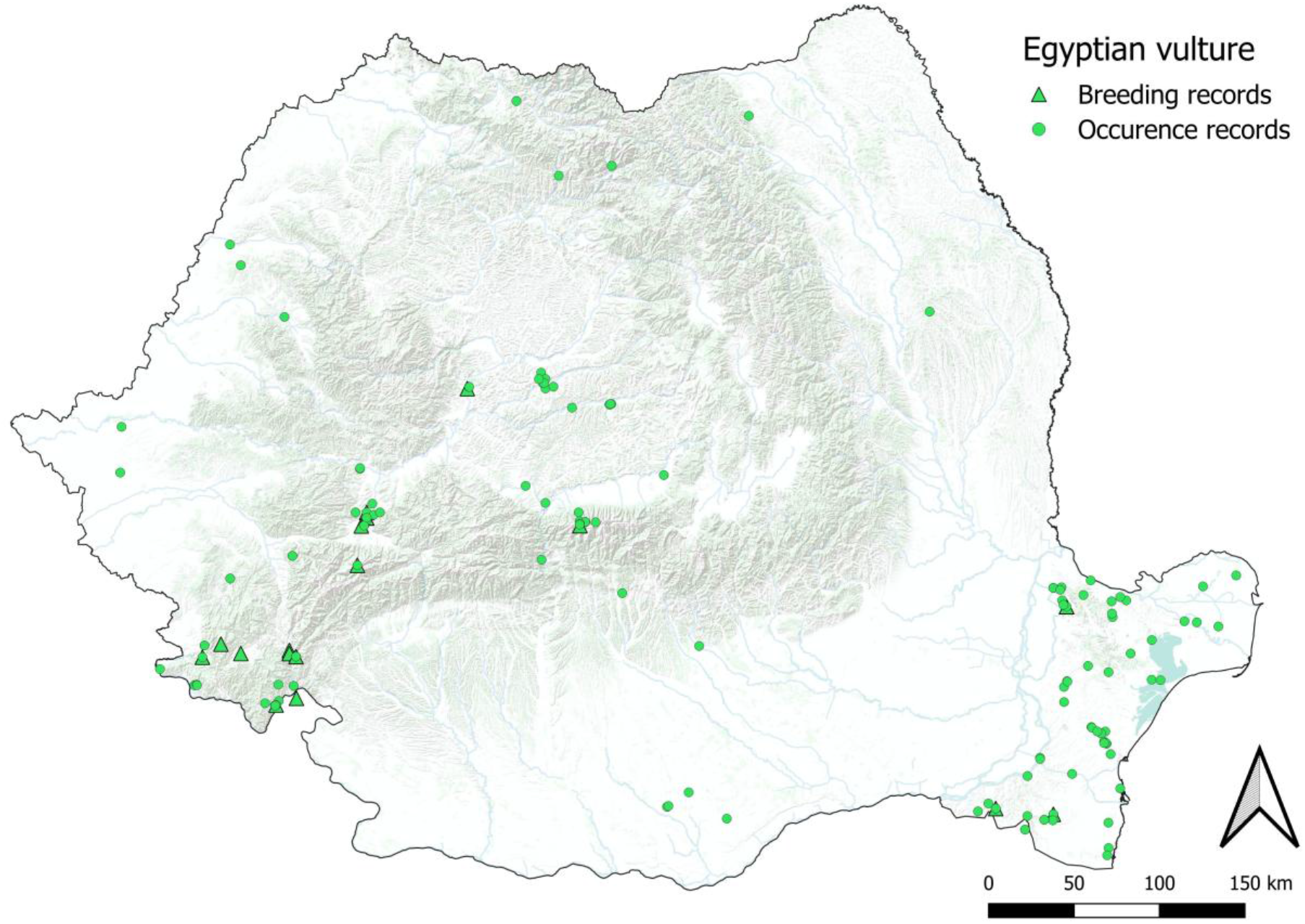
Distribution of *Neophron percnopterus* (Egyptian Vulture) in Romania. Green circles = all occurrence records (n = 173); green triangles = confirmed or probable breeding records (n = 23).

Twenty-three confirmed or probable breeding records span 1840–1966. Breeding in the Cazane area (Caraș-Severin) is documented from 1843 to at least 1962, making this the longest confirmed breeding locality in the database for any species. At Băneasa (Canaraua Fetei, Constanța County), breeding of a single pair was confirmed in 1966 (Cătuneanu et al. 1967; Paspaleva and Tălpeanu 1967) – the latest confirmed breeding record of any vulture species in Romania. Additional historical breeding occurred at Reea (Hunedoara, 1850), Dubova (Mehedinți, 1912) and Băneasa (Constanța, 1957).

### 3.6 *Gypaetus barbatus* (Bearded Vulture)

The database contains 315 records of Gypaetus barbatus, with 303 georeferenced (96.2%; Supplementary Material S2; Figure 7), the highest georeferencing rate among the four species. Czynk (1894) provided the first detailed monograph of the species in Transylvania, describing how in the 1860s–1870s Bearded Vultures regularly joined Griffon and Cinereous Vultures at the Brașov rendering yard, and noting that forest rangers routinely shot them without recognising the species, classifying them merely as “vultures” or “eagles.” This reflects the predominantly mountainous distribution of this species, where localities are generally precisely described. The temporal span is 1841–2016, and with a single exception (one record from 2016), all records pre-date 2000.

**Figure 7.**
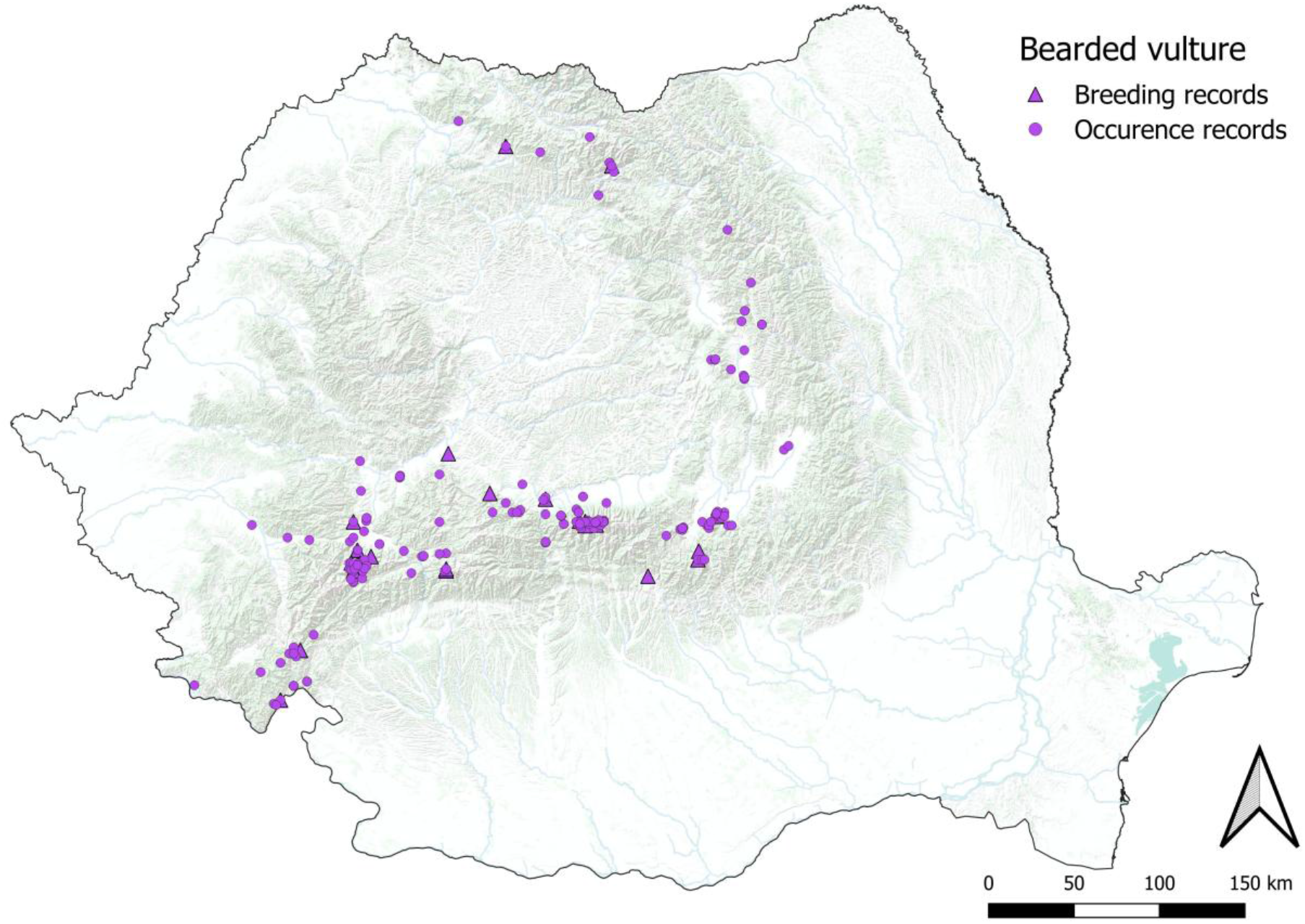
Distribution of *Gypaetus barbatus* (Bearded Vulture) in Romania. Purple circles = all occurrence records (n = 315); purple triangles = confirmed or probable breeding records (n = 40).

Forty confirmed or probable breeding records span 1841–1935. Hunedoara County (primarily the Retezat Mountains) yields the largest number of breeding records (n = 19) and the last confirmed breeding in the country: Stănuleți, Retezat Mountains, 1929 (Kamner 1928; Salmen 1980). A 1946 record from Tăul Custura Mare (guano and old nests found by geographer Dunăreanu) is considered unreliable. The Făgăraș Mountains (Sibiu and Argeș counties) represent the second major historical breeding area, with confirmed breeding at Negoiu Peak (Sibiu, 1889) and the Argeș flank of the Făgăraș range (1889). Additional historical breeding localities include Sebeș (Alba, 1873), Râu Mare in the Retezat (1873), and Suru Peak, Câineni on the Vâlcea flank of the Făgăraș (1927).

### 3.7 Spatial patterns and key sites

The Hunedoara–Sibiu–Brașov–Argeș corridor along the Southern Carpathian arc concentrates the highest density of historical breeding records for all four species, encompassing 32% of all confirmed breeding localities (Figure 8). Within this corridor, the Retezat Mountains (Hunedoara County) are the most important single locality, with breeding records for Griffon Vulture, Cinereous Vulture and Bearded Vulture. The Făgăraș Mountains (Sibiu, Brașov, Argeș, Vâlcea counties) hold confirmed breeding records for three of the four species (all except Egyptian Vulture). Argeș County holds 22 records across all four species, with Bearded Vulture contributing the largest share (n = 15), reflecting the extensive cliff systems of Cheile Argeșului (the upper Argeș valley) and adjacent Piatra Craiului area.

**Figure 8.**
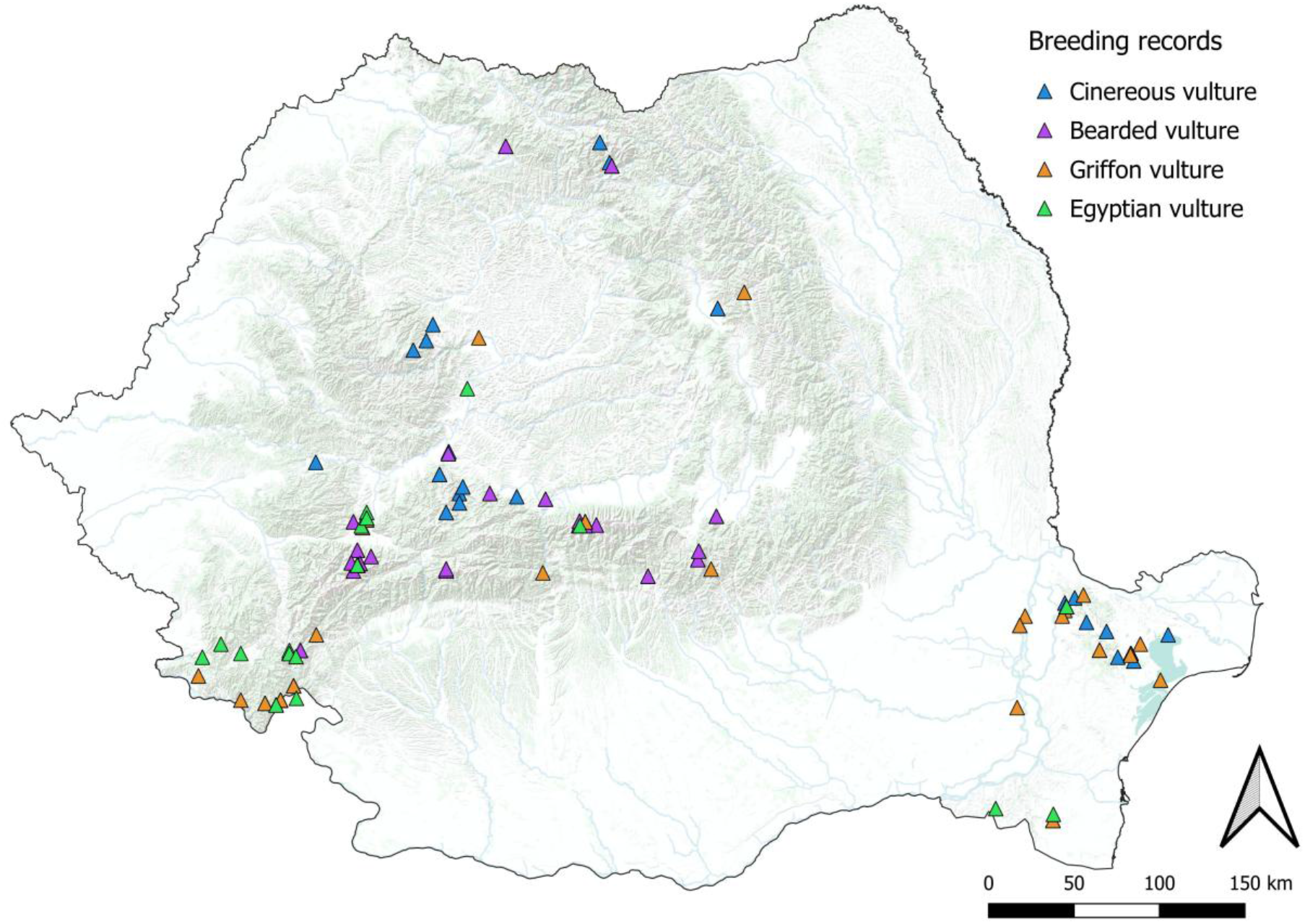
Distribution of confirmed and probable breeding records in Romania for the four vulture species. Triangles colour-coded by species.

Dobrogea represents the second major historical centre of vulture diversity, important primarily for Egyptian Vulture and Cinereous Vulture, but also yielding significant records of Griffon Vulture. The Babadag Forest (Tulcea) and Hagieni Forest (Constanța) are the most important Dobrogean localities, hosting last confirmed breeding for two species (Griffon Vulture 1929, Egyptian Vulture 1966). The Iron Gates (Cazane Gorge, Caraș-Severin/Mehedinți) is the third major cluster, with confirmed historical breeding of Griffon Vulture and Egyptian Vulture and the longest individual breeding site chronology in the database (Egyptian Vulture at Cazane, 1843–1962).

## 4. Discussion

The dataset spans more than two centuries of vulture records across present-day Romania, and three patterns stand out. All four species bred continuously across the Carpathian arc and Dobrogea throughout the nineteenth century. The entire breeding guild then collapsed within four decades, between the 1920s and 1960s. Non-breeding records have increased markedly since 2010, but this rebound reflects expanding Balkan source populations rather than recolonisation within Romania. We address in turn the causes and chronology of the collapse, the recent occurrence patterns, the conservation implications for guild recovery, and the limitations of the record.

### 4.1 Causes and chronology of breeding population collapse

The chronology of breeding collapse broadly mirrors European patterns (Donázar et al. 2016) but with pronounced region-specific characteristics. In Romania, direct persecution (shooting, strychnine and cyanide poisoning, and deliberate destruction of nest sites) is the most consistently cited cause in the historical literature. Griffon and Cinereous Vultures were widely regarded as livestock predators, and active eradication campaigns were conducted by agricultural authorities, particularly in Transylvania and the Carpathian foothills, from at least the 1870s onward; Comșia (1959) explicitly identified persecution as the principal factor in the local extinction of the Bearded Vulture from the Retezat. Specific episodes illustrate the scale: in January 1886 all ten Cinereous Vultures from a flock at Troschen perished after feeding on a strychnine-poisoned horse carcass (Csató, in Salmen 1980); a juvenile Bearded Vulture was found strychnine-poisoned below Hășmașul Mare in 1910 (Dobay, in Salmen 1980); and Tălpeanu (1967) reported the Egyptian Vulture as “constantly decreasing, probably always because of strychnine,” describing a specimen that fell dead in mid-flight at Băneasa in 1961. Intensive egg collection further contributed to the decline: the Sintenis brothers collected 377 Cinereous Vulture eggs from Babadag in three breeding seasons (Dombrowski 1912), and Dombrowski himself took 117 over his 21-year career in Romania (Cătuneanu 1973). The small Retezat and Făgăraș Bearded Vulture populations (probably fewer than five pairs at any point after 1900) were especially vulnerable to such additional mortality, and at least three eggs from Romanian breeding attempts are held in European museum collections.

The same threat complex (persecution combined with mass strychnine and cyanide poisoning campaigns targeting carnivores) drove Griffon Vulture to near-extinction in adjacent Bulgaria over the same period, with the population collapsing from over 1,000 pairs historically to near-zero by the 1960s before recovering under protection (Demerdzhiev et al. 2014). For Egyptian Vulture, the detailed documentation of extinction in south-eastern Bulgaria (Milchev and Georgiev 2014) reveals that shooting caused 50% of local extirpations, with addled eggs linked to livestock reduction reducing breeding success below replacement, a pattern closely paralleling the Romanian case.

The proximate role of persecution and poisoning operated against a backdrop of progressive food-base collapse, vividly articulated by Csató (1892, in Salmen 1980), who described the pre-railway era as “a golden age for the vultures,” when thousands of draft animals traversing Transylvania’s roads provided abundant carrion: “Now bad times have come for them. Dead cattle must be buried, therefore they get no food in the inhabited parts of the country and are confined to the high mountains.” Salmen (1980) attributed the Griffon Vulture’s retreat from the Transylvanian lowlands to late nineteenth-century sanitary regulations requiring burial of livestock carcasses. The twentieth-century transformation of Romanian livestock husbandry (abandonment of transhumant pastoralism, intensive stall-feeding, and from the late 1940s collectivisation of agriculture, which initially maintained large herds but dramatically reduced open-landscape carcass availability) progressively eroded the food base. Equivalent food-base mechanisms have been documented across Europe: declining sheep and goat livestock was the strongest predictor of Egyptian Vulture territory abandonment in Cantabrian Spain (Mateo-Tomás and Olea 2010), and in eastern Bulgaria Egyptian Vultures shifted from livestock carrion to wild prey after livestock reductions but were unable to compensate, going locally extinct (Milchev 2011). Salmen (1980) concluded that while nineteenth-century hunters decimated the Bearded Vulture, “it was strychnine in the twentieth century to which the last specimens of this majestic bird fell victim.” Dobay (1932) estimated only two Bearded Vulture pairs remained along the entire 600 km Carpathian arc from Orsova to Predeal; Csató had estimated 10–12 pairs for Retezat and Parâng alone in 1895 (Salmen 1980), illustrating a collapse from dozens of pairs to near-zero within four decades. By 1967, Tălpeanu concluded that the Griffon Vulture “no longer breeds in our country” and the Cinereous Vulture was known in Dobrogea “only by tradition” (Tălpeanu 1967); the last northern-Dobrogea records are 1954 (Griffon Vulture) and 1958 (Cinereous Vulture) (Cătuneanu 1973). The virtual elimination of free-ranging livestock and working horses from the Romanian countryside by the 1960s–1970s likely removed the last viable support system for any residual breeding populations.

The historical evidence assembled here carries direct implications for contemporary conservation planning. The three principal drivers of the Romanian vulture collapse (deliberate poisoning, direct shooting and food-base erosion) remain dominant threats to vulture populations across the Balkans and Mediterranean today. In reintroduced Griffon Vulture populations in central Italy, carbamate poisoning still accounts for 53% of recorded mortality (Posillico et al. 2023). In January 2026, six Cinereous Vultures and one Griffon Vulture were found dead from suspected illegal poisoning near Kotel in Bulgaria’s Stara Planina, the same region where the reintroduction programme had its greatest success. The victims included the first wild-hatched Cinereous Vulture recorded in Bulgaria in approximately 50 years (Vulture Conservation Foundation 2026). For Egyptian Vulture, population viability analysis of the Macedonian population (the nearest remnant to Romania’s historical Balkan range) predicts total extinction within 50 years under current mortality, with adult survival, not breeding productivity, as the critical parameter (Velevski et al. 2014). These data demonstrate that any Romanian recovery effort must be built on prior establishment of effective anti-poisoning protocols, power-line mitigation and legal enforcement against illegal killing.

### 4.2 Recent occurrence patterns and source population dynamics

The marked increase in non-breeding vulture records from 2010 onward reflects both the rapid growth of citizen-science recording in Romania and genuine changes in vagrancy patterns. The Balkan Griffon Vulture population has undergone a remarkable recovery, increasing to 445–565 pairs by 2019, although the breeding range has contracted to approximately half its 1980 extent and is now concentrated in three core subpopulations in Bulgaria, Serbia and Croatia (Dobrev et al. 2022). In Bulgaria, natural recovery in the Eastern Rhodopes (Demerdzhiev et al. 2014) was complemented by an ambitious reintroduction programme releasing over 275 individuals between 2010 and 2016, tripling the national breeding range (Stoynov et al. 2018); the reintroduced Eastern Balkan Mountains population comprised 23–25 breeding pairs by 2020 (Kmetova-Biro et al. 2021). The total Bulgarian population now exceeds 163 breeding pairs (Terraube et al. 2022), and GPS tracking has documented regular northward dispersal of juveniles and sub-adults into Romania, Serbia and Moldova (Stoychev et al. 2022). The pattern of recent Griffon Vulture records in our database, concentrated over the Southern Carpathians, with multiple records of soaring groups, is fully consistent with this dispersal dynamic, demonstrating that the Carpathian landscape already supports regular natural movement of the target species. The roost-prospecting behaviour documented in Bulgarian Griffon Vultures prior to colony establishment (Dobrev et al. 2020) suggests that some current Romanian observations may already represent site prospecting. Equivalent Central European trends were documented by Danko et al. (2013), who attributed influx years (2007, 2012) to a combination of food shortages in south-western populations and growing south-eastern source populations, with ringing recoveries directly tracing dispersing birds to Croatian and Serbian breeding colonies.

The dramatic increase in Cinereous Vulture records after 2020 (n = 61 in 2020–2025 vs. n = 7 in 2010–2019) is particularly striking. The predominantly immature age composition is consistent with the documented movement ecology of pre-adult Cinereous Vultures, in which juveniles consistently make the longest exploratory movements (Vágási et al. 2016; Tobajas et al. 2024), and mirrors the Eastern Rhodope pattern, where 72.7% of foraging Cinereous Vulture individuals from the Dadia colony were immature (Arkumarev et al. 2020). This likely reflects growth in regional Balkan source populations (most notably Serbia and the ongoing reintroduction in Bulgaria; Terraube et al. 2022), combined with increasingly systematic monitoring in Romania. Egyptian Vulture vagrants are now recorded annually in Constanța and Tulcea; these belong to the eastern Balkan flyway population, which migrates via Turkey and the Suez route to sub-Saharan wintering grounds (Oppel et al. 2022). Global MaxEnt habitat suitability modelling identifies south-eastern Romania as within the species’ predicted optimal range, with the principal habitat predictors (temperature seasonality, low precipitation in the coldest quarter) corresponding closely to the continental regime of Dobrogea (Panthi et al. 2021). The 2016 Bearded Vulture record is the first documented occurrence of this species in Romania since the 1960s and forms part of a growing body of evidence for long-distance dispersal of Alpine-origin birds reaching the Balkans (Frey et al. 2022); GPS tracking of Pyrenean Bearded Vultures has shown that non-territorial floaters use home ranges exceeding 10,000 km² (Margalida et al. 2016).

### 4.3 Conservation implications and priorities for recovery

Based on the spatial and temporal patterns documented in our database, nine areas emerge as historically well-supported candidate landscapes for vulture recovery in Romania: Domogled–Cerna Valley, Retezat Mountains, Almăj–Locvei Mountains, Făgăraș Piedmont, Cozia–Buila–Vânturarița Corridor, Trascău Mountains, Ceahlău Massif, Bicaz–Hășmaș Gorges and Nera–Beușnița Gorges. Each holds confirmed historical breeding records for at least two vulture species, and all lie within or adjacent to existing Special Protection Areas. Historical breeding evidence provides an essential first filter; final prioritisation requires contemporary habitat-suitability assessment, evaluation of current threat levels (poisoning, powerline collision, disturbance), food availability and assessment of social acceptance among local communities. The Făgăraș Mountains provide particularly strong support for the Griffon Vulture. The historical breeding range of the species in Romania extended from Cazane Gorge (Caraș-Severin, documented from 1835) through the Southern Carpathians, including the Argeș flank of the Făgăraș, with additional breeding records from Harghita and Prahova by the 1860s. The range also extended into Dobrogea, where breeding persisted until 1929. Mestecăneanu (2021) confirms Griffon Vulture as a frequent breeder in the eastern Făgăraș in 1920 (citing Jacobi 1984) and records Cinereous Vulture from the Făgăraș in the same year (Linția 1954). The Făgăraș–Argeș corridor was historically within the breeding range of the Bearded Vulture, with the latest reliable individual observations from the area dating to 1937–1938 (Călinescu 1938; Nania 1977; Băcescu 1961, all via Mestecăneanu 2021); breeding Cinereous Vulture is documented at the Sibiu/Făgăraș level in 1897, confirming that the cliffs, thermals and food base of this landscape historically supported the full guild. Recent records of soaring Griffon Vulture over the Făgăraș ridgeline in our database confirm that natural movement into the area is already occurring. Habitat-suitability modelling in Bulgaria identifies steep slopes, open landscapes and high livestock density as the primary predictors of Griffon Vulture breeding habitat (Dobrev and Popgeorgiev 2021), parameters amply present in the Făgăraș and Retezat candidate areas.

For Cinereous Vulture, the Retezat Mountains (Hunedoara) and Rodna Mountains (Bistrița-Năsăud) are the historically best-documented breeding strongholds. Both constitute logical priority areas for reintroduction feasibility assessment, subject to evaluation of current land use and human disturbance. The Retezat National Park, with its rugged cliffs, sub-alpine grasslands and existing large-raptor monitoring infrastructure, is particularly well-placed. Genetic evidence identifies Balkan and Iberian Cinereous Vulture as distinct evolutionary significant units (Poulakakis et al. 2008), indicating that Romanian source individuals should be drawn from Balkan-origin stock. The Spanish recovery demonstrates that population-level recovery is achievable within decades when legal protection, supplementary feeding and anti-poisoning measures are applied (Moreno-Opo and Margalida 2014).

For Bearded Vulture, the Retezat–Parâng–Făgăraș corridor emerges clearly as the primary historical range, with the Retezat the single most important locality in our database for the species (breeding documented 1841–1929). A Romanian reintroduction would ideally target the Retezat–Apuseni axis, consistent with the Alpine model of establishing populations in contiguous mountain ranges to enable eventual natural connectivity (Jenny et al. 2018). In May 2025 Bulgaria released the first three captive-bred Bearded Vultures into the wild at Sinite Kamani Nature Park (Sliven; Vulture Conservation Foundation 2025), and adopted its first National Action Plan for the Bearded Vulture (2025–2034), establishing a framework for recovery in the Eastern Balkans that could eventually extend to Romania’s Southern Carpathians.

For Egyptian Vulture, active reintroduction is probably premature given the continued and severe decline of the Mediterranean–Balkan metapopulation: across the Balkans the species has declined at annual rates λ = 0.920–0.951 (Velevski et al. 2015), with the breeding range fragmenting into six subpopulations and a 50-year extinction horizon predicted under current mortality (Velevski et al. 2014). The decline extends beyond the Balkans: in Armenia the population has fallen by ∼48% over 40 years to just 39–43 breeding pairs (Aghababyan et al. 2026). Adult mortality from poisoning, electrocution and food shortage, not breeding failure, is the critical demographic parameter. However, the historical concentration of records in Dobrogea and the Iron Gates, combined with continued vagrant occurrences, suggests natural recolonisation remains possible if the Balkan population recovers. Global MaxEnt modelling places Dobrogea within the predicted core suitable range (Panthi et al. 2021), meaning habitat, not source-population pressure, is the current limit. Habitat management in Dobrogean SPAs (Băneasa–Canaraua Fetei, Hagieni Forest, Dobrogea Gorges) and the lower Danube gorge to maintain suitable nesting cliffs and increase carrion availability should be a medium-term priority.

Community acceptance is a critical, often underestimated, component of reintroduction feasibility. The only published baseline on Romanian public attitudes (Mertens et al. 2005), based on 292 interviews across nine communes within the Retezat National Park, reported broadly encouraging results: 88.7% of respondents stated they would support a Griffon Vulture reintroduction, 79.8% would be pleased to see vultures in their home area, and only 5.8% expressed fear of reintroduction. The same survey, however, revealed substantial knowledge deficits: most respondents incorrectly believed vultures to be active predators that kill livestock — beliefs that more accurately describe eagles, with which vultures are frequently confused. Targeted public education clearly differentiating vultures from predatory raptors should therefore be considered an integral component of any future reintroduction programme rather than a secondary activity. These figures should also be re-evaluated against the current social context, given the substantial demographic, economic and cultural changes in rural Romania since 2005.

Vulture reintroduction is now actively underway in Romania. Foundation Conservation Carpathia (FCC), in partnership with the Vulture Conservation Foundation (VCF) and Milvus Group, initiated the first Griffon Vulture reintroduction in Romanian history: 25 individuals sourced from Spain arrived at an acclimatisation aviary near Rucăr (Argeș County, 1,150 m a.s.l.) in March 2026, with release into the Făgăraș Mountains planned after a six-month acclimatisation period. A second Griffon Vulture project, led by Rewilding Romania at Obârșia Cloșani in the Domogled–Cerna Valley National Park, is in the aviary construction phase, with first individuals scheduled for September 2026. Both target landscapes of confirmed historical breeding and are supported by area-specific feasibility assessments: for the Făgăraș massif (Nagy et al. 2024) and for the Retezat National Park (Kelemen and Mertens 2006); the Domogled–Cerna candidate area lies within the same south-western Carpathian massif. The historical baseline assembled here can serve as a reference for identifying additional suitable areas for future vulture recovery in Romania.

More broadly, the Romanian case illustrates a general principle: in countries where vulture populations were lost before modern monitoring infrastructure was established, the absence of systematic baseline data risks creating a shifting baseline against which recovery targets are set too conservatively, release sites are selected on incomplete information, and the spatial extent of formerly suitable habitat is underestimated. The present database demonstrates that, even in data-poor regions, the integration of museum collections, multilingual historical literature and citizen-science records can recover sufficient information to guide the reassembly of functionally extinct species guilds.

Despite its breadth, the dataset remains uneven in space and time. Transylvania and Dobrogea are disproportionately represented because of the intensity of historical ornithological activity and museum collecting in these regions, whereas Muntenia, Oltenia, Moldova and Bukovina remain under-documented. Consequently, the last breeding dates reported here should be interpreted as last documented breeding records rather than absolute extinction dates. For the Cinereous Vulture, the May 1942 breeding record at Rodna (Salmen 1980) is the most reliable breeding endpoint, the often-cited “c. 1964” (Baumgart 1974a, citing Klemm 1966) being a tertiary claim without primary verification. Sporadic Egyptian Vulture breeding in the Iron Gates or Dobrogean gorges after 1966 also cannot be excluded on current evidence. Additional work in regional museums, historical archives and under-surveyed Romanian ornithological literature, especially for the period 1950–2009, may refine the chronology of decline and reveal further records, particularly from the Eastern Carpathians and Dobrogea, for which even the earliest systematic source for the region (Nordmann 1840) is fragmentary.

## 5. Conclusions

Romania holds a position of exceptional importance in European vulture conservation: it is one of the few EU member states in which all four Old World vulture species – *Gyps fulvus*, *Aegypius monachus*, *Neophron percnopterus* and *Gypaetus barbatus* – historically maintained confirmed breeding populations. Our database of 1,136 records spanning 1818–2025 constitutes the most comprehensive historical baseline yet assembled for these species in Romania. It documents widespread nineteenth-century breeding across the Carpathian arc and Dobrogea, followed by near-total collapse between the 1920s and 1960s driven by direct persecution, poisoning and agrarian transformation. The staggered sequence of breeding extinction – G. fulvus 1929, G. barbatus 1929, A. monachus 1942, N. percnopterus 1966 – reflects differences in ecology, vulnerability to persecution and the extent of each species’ breeding range.

The Făgăraș and Retezat Mountains emerge as the most important historical breeding strongholds for three of the four species, validating the choice of the Făgăraș Mountains as the site of Romania’s first Griffon Vulture reintroduction programme. The marked and continuing increase in non-breeding records of Griffon Vulture and Cinereous Vulture after 2010 confirms growing ecological connectivity between Romanian habitats and expanding Balkan source populations, providing favourable conditions for both reintroduction support and eventual natural recolonisation. Future work should prioritise archival and museum searches in under-documented regions, targeted reassessment of historical breeding areas, and integration of the present baseline with contemporary habitat suitability, threat and social-acceptance analyses. Together, these steps can help guide the evidence-based reassembly of Romania’s lost vulture guild.

## Acknowledgements

Special thanks go to Cristi Domșa, who gave us access to the database of the Romanian Ornithological Society, and to József Szabó and Szilárd Daróczi for access to the Rombird database. We are also thankful to the Milvus Group Bird and Nature Protection Association for making vulture observations stored in the openbirdmaps database available to us. We thank Natalia Irimuș and Ágnes Brem for helping us access historical literature held by the Central University Library of Babeș-Bolyai University. Thanks to Andreea Cătălina Drăghici for helping in scientific literature search. We also thank Cerchia Corneliu (Vasile Pârvan Museum, Bârlad) and Ioana Vanesa Duceac (Muzeul de Zoologie, Facultatea de Biologie, Universitatea „Alexandru Ioan Cuza” din Iași) and Medeea Hălici (Muzeul de Științele Naturii, Dorohoi), Iulian Laurențiu Mazilu (Muzeul Vrancei, Focșani), Sike Tamás (Muzeul Județean Satu Mare), Silviu Țicu (Muzeul Național Brukenthal, Sibiu) for providing information about vulture species in their collections. Thanks to Tamás Papp, Szilárd Daróczi, and Dragomir-Cosmin David for the comments to the preprint version. Special thanks to Osváth-Ferencz Márta for her support.

## Author Contributions

Gergely Osváth: Conceptualization, Methodology, Project administration, Data curation, Investigation, Visualization, Writing the first version of the manuscript.

Avar-Lehel Dénes: Conceptualization, Methodology, Data curation, Validation, Writing – review and editing.

Zsolt Kovács: Investigation, Data curation, Writing – review and editing.

Alexandru-Cătălin Birău: Investigation, Data curation, Writing – review and editing.

Edgár Papp: Visualization, Data curation, Writing – review and editing.

Gabriella-Veronka Jákó: Investigation, Data curation, Writing – review and editing.

Róbert Zeitz: Conceptualization, Methodology, Investigation, Resources, Writing – review and editing.

All authors read, edited, and approved the final version of the manuscript.

## Data availability statement

The complete occurrence database is provided as Supplementary Material S2, the detailed methodology as Supplementary Material S1, and the interactive visualisation as Supplementary Material S3. Upon acceptance, these materials will be archived in a publicly accessible repository, with a DOI provided in the final version.

## Supplementary materials

*Supplementary Materials S1–S3 are available with the online version of this article*.

Supplementary Material S1. Methods and references. Detailed account of materials and methods, including study area description, source categories, database structure, quality control protocols, data analysis workflow, and the complete bibliography of all consulted sources (provided as a .docx file).

Supplementary Material S2. Complete occurrence database (1,136 records, 1818–2025). Fields: species, location, county, region, latitude, longitude, date, year, number of individuals, record type, status, certainty, observer, age, sex, source, citation, notes. Provided as an Excel file (.xlsx).

Supplementary Material S3. Interactive offline-capable data visualisation of the complete database.

